# Microstructure Diffusion Scalar Measures from Reduced MRI Acquisitions

**DOI:** 10.1101/772897

**Authors:** Santiago Aja-Fernández, Rodrigo de Luis-García, Maryam Afzali, Malwina Molendowska, Tomasz Pieciak, Antonio Tristán-Vega

## Abstract

In diffusion MRI, the Ensemble Average diffusion Propagator (EAP) provides relevant microstructural information and meaningful descriptive maps of the white matter previously obscured by traditional techniques like the Diffusion Tensor. The direct estimation of the EAP, however, requires a dense sampling of the Cartesian q-space. Due to the huge amount of samples needed for an accurate reconstruction, more efficient alternative techniques have been proposed in the last decade. Even so, all of them imply acquiring a large number of diffusion gradients with different b-values. In order to use the EAP in practical studies, scalar measures must be directly derived, being the most common the return-to-origin probability (RTOP) and the return-to-plane and return-to-axis probabilities (RTPP, RTAP).

In this work, we propose the so-called “Apparent Measures Using Reduced Acquisitions” (AMURA) to drastically reduce the number of samples needed for the estimation of diffusion properties. AMURA avoids the calculation of the whole EAP by assuming the diffusion anisotropy is roughly independent from the radial direction. With such an assumption, and as opposed to common multi-shell procedures based on iterative optimization, we achieve closed-form expressions for the measures using information from one single shell. This way, the new methodology remains compatible with standard acquisition protocols commonly used for HARDI (based on just one b-value). We report extensive results showing the potential of AMURA to reveal microstructural properties of the tissues compared to state of the art EAP estimators, and is well above that of Diffusion Tensor techniques. At the same time, the closed forms provided for RTOP, RTPP, and RTAP-like magnitudes make AMURA both computationally efficient and robust.

## 1. Introduction

Under the name of Diffusion Magnetic Resonance Imaging (DMRI) we gather a set of diverse MRI imaging techniques with the ability of extracting *in vivo* relevant information regarding the random, anisotropic diffusion of water molecules that underlay the structured nature of different living tissues. It has attracted an extraordinary interest among the scientific community over recent years due to the relationships found between a number of neurological and neurosurgical pathologies and alterations in the white matter as revealed by an increasing number of DMRI studies (Rovaris and Filippi, 2007; Bester et al., 2015; Pasternak et al., 2015).

The most relevant feature of DMRI is its ability to measure directional variance, i.e. anisotropy. In the beginning of the 2000s, diffusion tensor MRI (Basser and Pierpaoli, 1996, DT-MRI) gained huge popularity in white matter studies, not only among technical researchers but also among clinical partners, to the point that even nowadays most of the research studies involving DMRI focus on the diffusion tensor (DT). By using a simple Gaussian regressor, the anisotropy of the tissues is actually probed by acquiring as few as 20 to 60 images, which is acceptable in clinical practice. DT-MRI brought to light one of the most common problems in DMRI techniques: in order to carry out clinical studies, the information given by the selected diffusion analysis method must be translated into some scalar measures that describe different features of the diffusion within every voxel. That way, measures like the Fractional Anisotropy (FA), the Axial and Radial Diffusivity (AD, RD) or the Mean Diffusivity (MD) were defined (Westin et al., 2002). Even at the early stages of DT-MRI, it was clear that the Gaussian assumption had important limitations. It provided a useful tool allowing clinical studies, but the underlying diffusion processes were not accurately described because of the over-simplified fitting, so that more evolved techniques with more degrees-of-freedom naturally arose, such as High Angular Resolution Diffusion Imaging (Tuch et al., 2003; Tristán-Vega et al., 2009; Özarslan et al., 2006, HARDI) or Diffusion Kurtosis Imaging (Jensen et al., 2005, DKI). It seems obvious that more degrees-of-freedom require more diffusion images to be acquired, but the requirement of an accurate angular resolution also implies the need for a finer angular contrast, which translates in the need for stronger gradients to probe diffusion, i.e., higher b-values (LeBihan, 1991).

The trend over the last decade has consisted in acquiring a large number of diffusion-weighted images distributed over several shells together with moderate-to-high b-values to estimate more advanced diffusion descriptors, such as the Ensemble Average diffusion Propagator (Wedeen et al., 2005; Özarslan et al., 2013, EAP). This leads to a completely model-free, non parametric approach for diffusion that can accurately describe most of the relevant phenomena associated to diffusion.

The most straightforward way of estimating the EAP is Diffusion Spectrum Imaging (Wedeen et al., 2005, DSI), that relies on the dense sampling of the q-space for discrete Fourier transformation. Hence, it requires a huge number of images to avoid aliasing artifacts and attain a decent accuracy, which makes it not so appealing in practice. As a consequence, alternative techniques aim to reconstruct the EAP from reduced samplings of the q-space, most of them related to the recent advances in compressed sensing and sparse reconstruction (Landman et al., 2012; Merlet and Deriche, 2010). In practice, some multishell reconstruction techniques may be used to compute the EAP, typically as a superposition of the integrals analytically computed for each basis function. Some of the most prominent methods are *Hybrid Diffusion Imaging* (HYDI) by Wu and Alexander (2007), the *multiple q-shell diffusion propagator imaging* (mq-DPI) by Descoteaux et al. (2009, 2011), the *Bessel Fourier Orientation Reconstruction* (BFOR) by Hosseinbor et al. (2013), the *directional radial basis functions* (RBFs), recently proposed by Ning et al. (2015), or the *Simple Harmonic Oscillator Based Reconstruction and Estimation* by (Özarslan et al., 2008, SHORE). More recently, the *Mean Apparent Propagator MRI* (MAP-MRI) proposed by Özarslan et al. (2013) and its improved version, the so-called *Laplacian-regularized MAP-MRI* (Fick et al., 2016b, MAPL), have gained interest among the community due to the compelling results demonstrated in several clinical trials (Avram et al., 2016).

There is no doubt the EAP provides rich and valuable anatomical information about the diffusion process. However, such amount of information may result overwhelming and difficult to integrate within clinical studies. This pitfall is usually circumvented by defining scalar measures directly related to the characteristics of diffusion acting as biomarkers candidates (such as diffusivity, anisotropy, intra-cellular vs. extra-cellular water movement, etcetera). Some late examples are the probability of zero displacement (or return-to-origin probability, RTOP), the mean-squared displacement (MSD), the q-space inverse variance (QIV), or the return-to-plane and return-to-axis probabilities (RTPP, RTAP) (Hosseinbor et al., 2012; Wu et al., 2008; Ning et al., 2015).

Although the use of these measures is not generalized among the clinical community, there is a growing interest on the exploration of their potential clinical applicability. To date, the relevance of scalar descriptors of the brain microstructure has been proved on both *ex vivo* (Daianu et al., 2015; Özarslan et al., 2013; Fick et al., 2016a) and *in vivo* studies of healthy and diseased subjects (Brusini et al., 2015; Hosseinbor et al., 2012; Avram et al., 2016; Brusini et al., 2016; Boscolo Galazzo et al., 2018; Zucchelli et al., 2016). In particular, RTOP has also demonstrated to be a better indicator for cellularity and diffusion restrictions than the DTI-related mean diffusivity (MD) (Avram et al., 2016) and, together with MSD, a proper measure for the assessment of myelination (Alimi et al., 2018). These results were later confirmed by Boscolo Galazzo et al. (2018), where the authors reinforced the hypothesis on RTOP to have greater specificity to reflect cellularity and restricted diffusion.

The obvious drawback of this methodology is the need of acquiring very large data sets with many q-space samples in different shells (some of them with very large b-values, which implies an additional problem due to noise, eddy currents, non-linear effects, etcetera). Even when sophisticated non-linear techniques based on compressed sensing are used, the number of gradient images to be acquired vastly exceeds that needed for single-shell protocols like DTI or HARDI. This is clearly a practical limitation: a large number of samples goes together with longer scanning times, subject movement, and patient discomfort that make them unfit for clinical practice and for many clinical studies. Besides, some methods require b-values that cannot be acquired by some commercial scanners.

The present paper delves into the previously introduced hypothesis that scalar measures such as RTOP, RTPP, or RTAP are intrinsically tied up to the computation of the whole, model-free EAP or, on the contrary, a constrained model for radial diffusion may reveal analogous information using simpler protocols. To that end, we have first reformulated all these indices for single-shell acquisitions based on different diffusion models, so that the corresponding scalar measures are compatible with more standard acquisition protocols like those used in HARDI or DKI, i.e., with faster acquisitions based on heavily reduced data sets. The indices calculated with this new methodology are extensively tested against EAP-based indices to check if both sets are actually comparable.

## 2. Background

### 2.1. The Diffusion signal

The EAP, *P*(**R**), is the three dimensional Probability Density Function (PDF) of the water molecules inside a voxel moving an effective distance **R** in an effective time *τ*. It is related to the normalized magnitude image provided by the MRI scanner, *E*(**q**), by the Fourier transform (Callaghan et al., 1988):

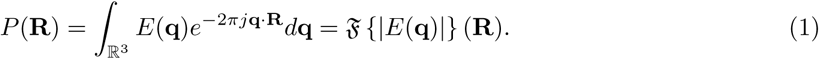

The inference of exact information on the **R**–space would require the sampling of the whole **q**–space to use the Fourier relationship between both spaces.

In order to obtain a closed-form analytical solution from a reduced number of acquired images, a model for the diffusion behavior must be adopted. The most common techniques rely on the assumption of a Gaussian diffusion profile and a steady state regime of the diffusion process that yields to the well-known Diffusion Tensor (DT) approach. Alternatively, a more general expression for *E*(**q**) can be used (Özarslan et al., 2006):

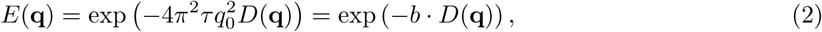

where the positive function *D*(**q**) = *D*(*q*_0_, *θ, ϕ*) *>* 0 is the Apparent Diffusion Coefficient (ADC), *b* = 4*π*^2^*τ* ‖**q**‖ ^2^ is the so-called b-value and *q*_0_ = ‖**q**‖, and *θ, ϕ* are the angular coordinates in the spherical system. According to Basser (2002), in the mammalian brain, this mono-exponential model is predominant for values of b up to 2000 s/mm^2^ and it can be extended to higher values (up to 3000 s/mm^2^) if appropriate multicompartment models of diffusion are used.

### 2.2. Advanced diffusion scalar measures

Although the EAP provides the global information about the diffusion in every voxel of the brain, that information must be properly translated to be used in clinical trials or to study the features of particular tissues. Regardless of the method used to estimate the EAP, it must provide a set of metrics to inspect the changes of complex brain microstructures, e.g., multiple compartments or restricted diffusion. Some of the most relevant measures usually derived from the EAP are:

1. **Return-to-origin probability (RTOP):** also known as *probability of zero displacement*, it is related to the probability density of water molecules that minimally diffuse within the diffusion time *τ*. It is known to provide relevant information about the white matter structure (Assaf et al., 2000; Hosseinbor et al., 2012; Wu et al., 2008), and has demonstrated to be a better indicator for cellularity and diffusion restrictions than the DTI-related mean diffusivity (MD) (Avram et al., 2016). It is defined as the value of *P*(**R**) at the origin, related to the volume of the signal *E*(**q**):

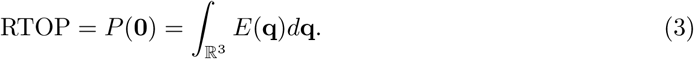
2. **Return-to-plane probability (RTPP)**, defined as:

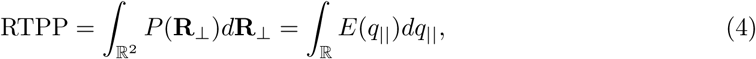

where *q*_‖_ denotes the direction of maximal diffusion. It is known to be a good indicator of restrictive barriers in the axial orientation, and it is related to the mean pore length (Özarslan et al., 2013).
3. **Return-to-axis probability (RTAP)**, defined as:

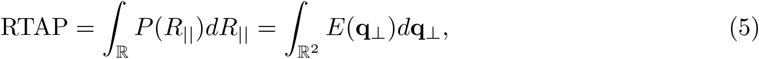

where **q**_⊥_ is the set of directions perpendicular to **q**_‖_ (the one with maximal diffusion). It is also a directional scalar index and an indicator of restrictive barriers in the radial orientation. According to Özarslan et al. (2013), RTPP and RTAP values can be seen as the *decomposition* of the RTOP values into two components, parallel and perpendicular to the maximum diffusion.

## 3. Methods

### 3.1. Diffusion measures from single shell acquisitions

The estimation of a given magnitude is always a trade-off between the available data and the complexity of the model. In this case, we consider a single shell acquisition compatible with HARDI: moderated-to-high b-value (ranging from 2000 to 3000 s/mm^2^) and moderated-to-large number of gradients. Since the amount of data is reduced, we are forced to assume a restricted diffusion model consistent with single-shell acquisitions: the ADC will be roughly independent from the radial direction within the range of b-values probed, so that *D*(**q**) = *D*(*θ, ϕ*). This way eq. (2) becomes:

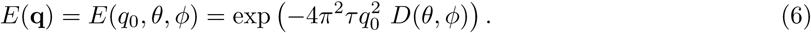

Note that, even when *D*(**q**) does not depend on *q*_0_, *E*(**q**) does. Once we cast this model into the scalar measures previously defined, we can simplify their formulations accordingly:

1. **RTOP:** By using the simplification in eq. (6), we can write eq. (3) in spherical coordinates and integrate with respect to the radial component *q*_0_:

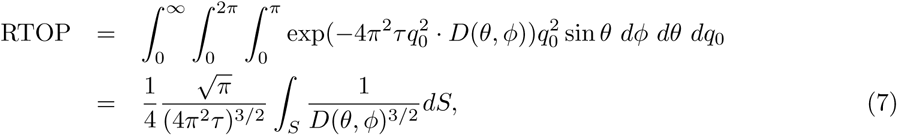

where *∫_S_* denotes the integral in the surface of a sphere *S* of radius one. This way, the integration in the whole **q**-space in eq. (3) reduces to the integration on the surface of a single shell.
2. **RTPP:** The diffusion signal *D*(**q**) in the maximum diffusion direction is given by *D*(*r*_0_), with *r*_0_ = *q*_‖_. Since that direction does not depend on *q*_0_, we can integrate with respect to the radial component:

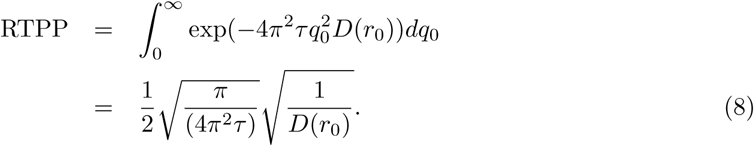
3. **RTAP:** Let *θ*′ be the angle that parametrizes the equator normal to the maximum diffusion direction and *D*(*θ*′) the diffusion signal at that equator. Once more, *D*(*θ*′) does not depend on the radial component and the integral can be solved:

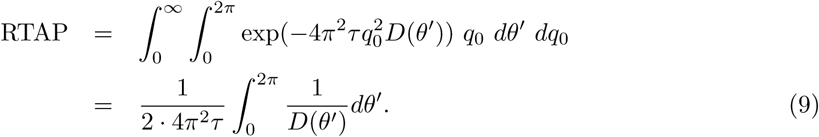 The original integral reduces to the line integral of a function in a plane perpendicular to the maximum diffusion direction.

Although the mono-exponential assumption in eq. (6) may seem restrictive, it has been successfully adopted before for single-shell, HARDI models to accurately describe several predominant diffusion directions within the imaged voxel (Descoteaux et al., 2006; Frank, 2002; Özarslan et al., 2006; Tristán-Vega et al., 2009). Moreover, it allows to get rid of the dense sampling required by the original formulations of RTOP, RTPP, and RTAP, as long as the volumetric integrals over the whole q-space are replaced by surface integrals over one single shell.

On the other hand, the mono-exponential model will roughly hold only within a limited range around the measured b-value, but diffusion features will radically differ for very different b-values. Consequently, it seems unlikly that the measures will show the exact same values for the whole range of b. For this reason, the measures derived this way must be seen as *apparent values* at a given b-value, related to the original ones but dependent on the selected shell. In what follows, they will be referred to as **Apparent Measures Using Reduced Acquisitions (AMURA)**.

### 3.2. Practical Implementation

Different methods can be applied in order to numerically calculate the AMURA expressions given in the previous section from the available data. In what follows, we propose a specific numerical procedure that ensures a robust calculation of the measures:

1. **RTOP:** the integration on the surface of the unit sphere *S* from a limited number of samples is calculated using Spherical Harmonics (SH):

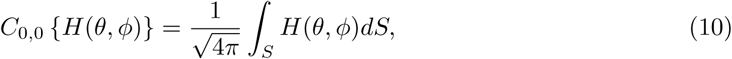

where *C*_0,0_ *{H*(*θ, ϕ*) *}* is the 0 *−* th order coefficient of the SH fitted to the signal *H*(*θ, ϕ*). This way, the RTOP becomes:

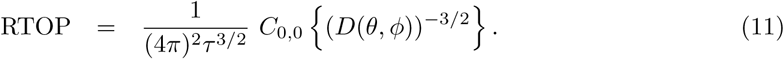
2. **RTPP:** The value of RTPP previously defined in eq. (8) depends on *D*(*r*_0_), the ADC evaluated at the direction of maximum diffusion. In order to avoid the variability that a maximum operator may introduce, we calculate the index over a regularized version of *D*(*θ, ϕ*). Let us call *D*_SH_(*θ, ϕ*) a version of the original diffusion signal regularized using SH. Then, we can write the RTPP as

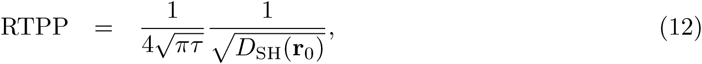

where **r**_0_ denotes the maximum diffusion direction.
3. **RTAP:** The value of 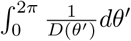 is the line integral of the function 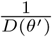 in a plane normal to the maximum diffusion direction. That integral is usually computed as the Funk-Radon transform (FRT) of such function, *𝒢{.}*, after Tuch (2004):

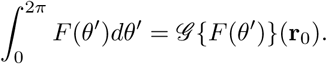 Therefore we can calculate eq. (9) as

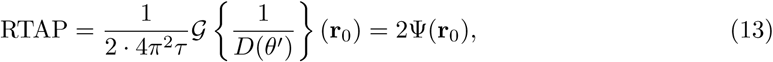

where Ψ(**r**) is the pQ-Balls whose definition and SH-based numerics were stated by Canales-Rodríguez et al. (2009); Tristan-Vega et al. (2010), and **r**_0_ is the direction of maximum diffusion.

An overview of AMURA, together with the specific numerical implementation of each magnitude, is presented in Table 1.

**Table 1:**
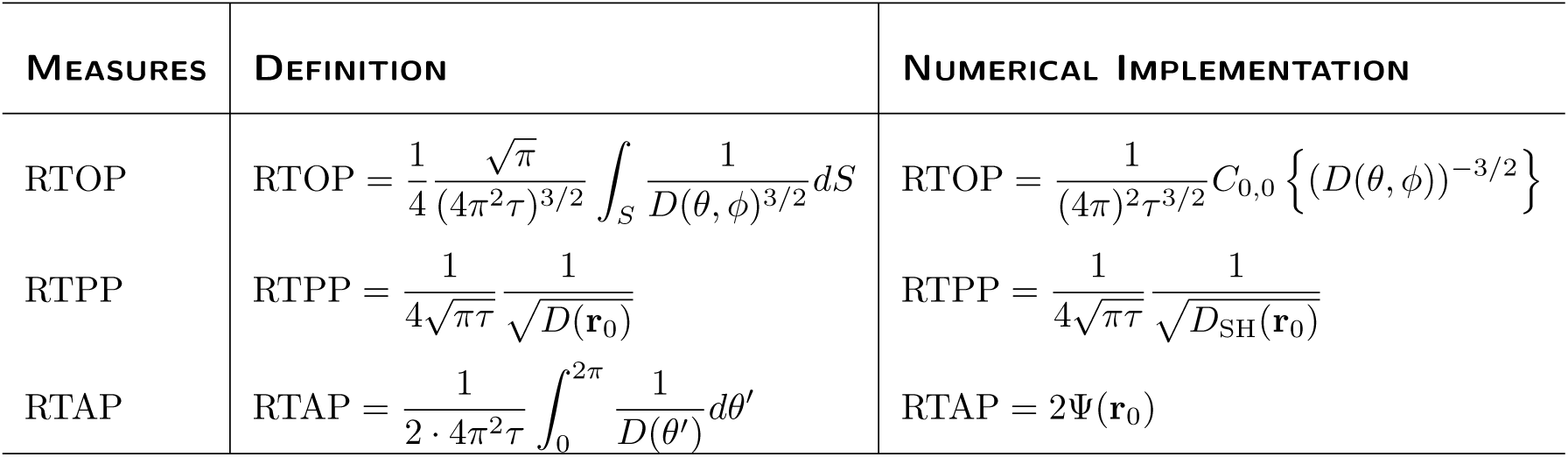
Survey of the q-space measures gathered by AMURA, along with their specific numerical implementations.

## 4. Results

### 4.1. Setting-up of the experiments

The computation of AMURA, as described above, relies on a number of parameters. The orientation functions computed, *E*(**q**), *D*(**q**), and Ψ(**r**), are represented in the basis of SH, whose coefficients are fitted via regularized Least Squares (LS) as described by Descoteaux et al. (2007). SH expansions of even orders up to 6 are considered in all cases, with a Tikhonov regularization parameter *λ* = 0.006. RTAP is computed from pQ-Balls as described by Tristan-Vega et al. (2010).

For the sake of replicability, only public data sets are considered for the experiments:

1. From the Human Connectome Project (HCP)^1^, five volumes were chosen: MGH 1007, MGH 1010, MGH 1016, MGH 1018 and MGH 1019, acquired in a Siemens 3T Connectome scanner with 4 different shells at *b* = [1000, 3000, 5000, 10000] s/mm^2^, with [64, 64, 128, 256] gradient directions each, in-plane resolution 1.5 mm^2^, and slice thickness 1.5 mm. Acquisition parameters are TE=57 ms and TR=8800 ms. The acquisition included 40 different baselines that were averaged to improve their SNR^2^.
2. From the Public Parkinson’s disease database (PPD)^3^, 53 subjects from a cross-sectional Parkinson’s disease (PD) study comprising 27 patients together with 26 age, sex, and education-matched control subjects. Data were acquired on a 3T head-only MR scanner (Magnetom Allegra, Siemens Medical Solutions, Erlangen, Germany) operated with an 8-channel head coil. Diffusion-weighted (DW) images were acquired with a twice-refocused, spin-echo sequence with EPI readout at two distinct b-values *b* = [1000, 2500] s/mm^2^, and along 120 evenly spaced encoding gradients. For the purposes of motion correction, 22 unweighted (b = 0) volumes, interleaved with the DW images, were acquired. Acquisition parameters are TR=6800 ms, TE=91 ms, and FOV=211 mm^2^, no parallel imaging and 6/8 partial Fourier were used. More information can be found in Ziegler et al. (2014).

### 4.2. Comparison with multi-shell EAP estimators

The working hypothesis in the present paper is that measures obtained with AMURA are closely related to the same measures derived using multi-shell-derived approaches. Although different assumptions are considered, it is expected that the different approaches assess similar diffusion properties. To test this hypothesis, the correlation between the measures computed with the different methods is estimated.

Three different EAP estimators have been selected for this experiment: the RBFs with constrained *ℓ*_2_ regularization and standard parameters suggested by the authors (Ning et al., 2015), the MAP-MRI with anisotropic basis and radial order 6 (Özarslan et al., 2013), and the MAPL with anisotropic basis, radial order 8, and regularization weighting *λ* = 0.2 (Fick et al., 2016b). This selection has deliberately excluded methods requiring a dense or Cartesian sampling of the q-space, since the HCP dataset includes just 4-shell acquisitions. Besides, the code for all the selected methods has been made publicly available.

To reduce the amount of data and keep a reasonable complexity (and reasonable computing times), only three different slices from each selected HCP volume were selected, see Table 2. The measures are calculated for RBF, MAP-MRI and MAPL using either 3 shells (b= {1000, 3000, 5000}), or 2 shells (b= {1000, 3000}). Their outermost shell (b=10000) has been discarded for this experiment, since such a high b-value will give rise to very different diffusion properties. AMURA is calculated using one single shell for two different b-values, b=3000 or b=5000. A white matter mask is used to reject voxels with FA*<* 0.2 for the computation of both the raw measures in Fig. 1 and the **correlations** in Table 3.

**Table 2:**
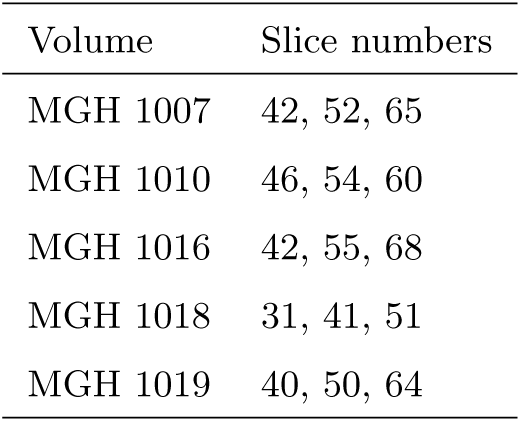
Selected slices from each diffusion volume from the HCP.

| Volume | Slice numbers |
| --- | --- |
| MGH 1007 | 42, 52, 65 |
| MGH 1010 | 46, 54, 60 |
| MGH 1016 | 42, 55, 68 |
| MGH 1018 | 31, 41, 51 |
| MGH 1019 | 40, 50, 64 |

**Table 3:**
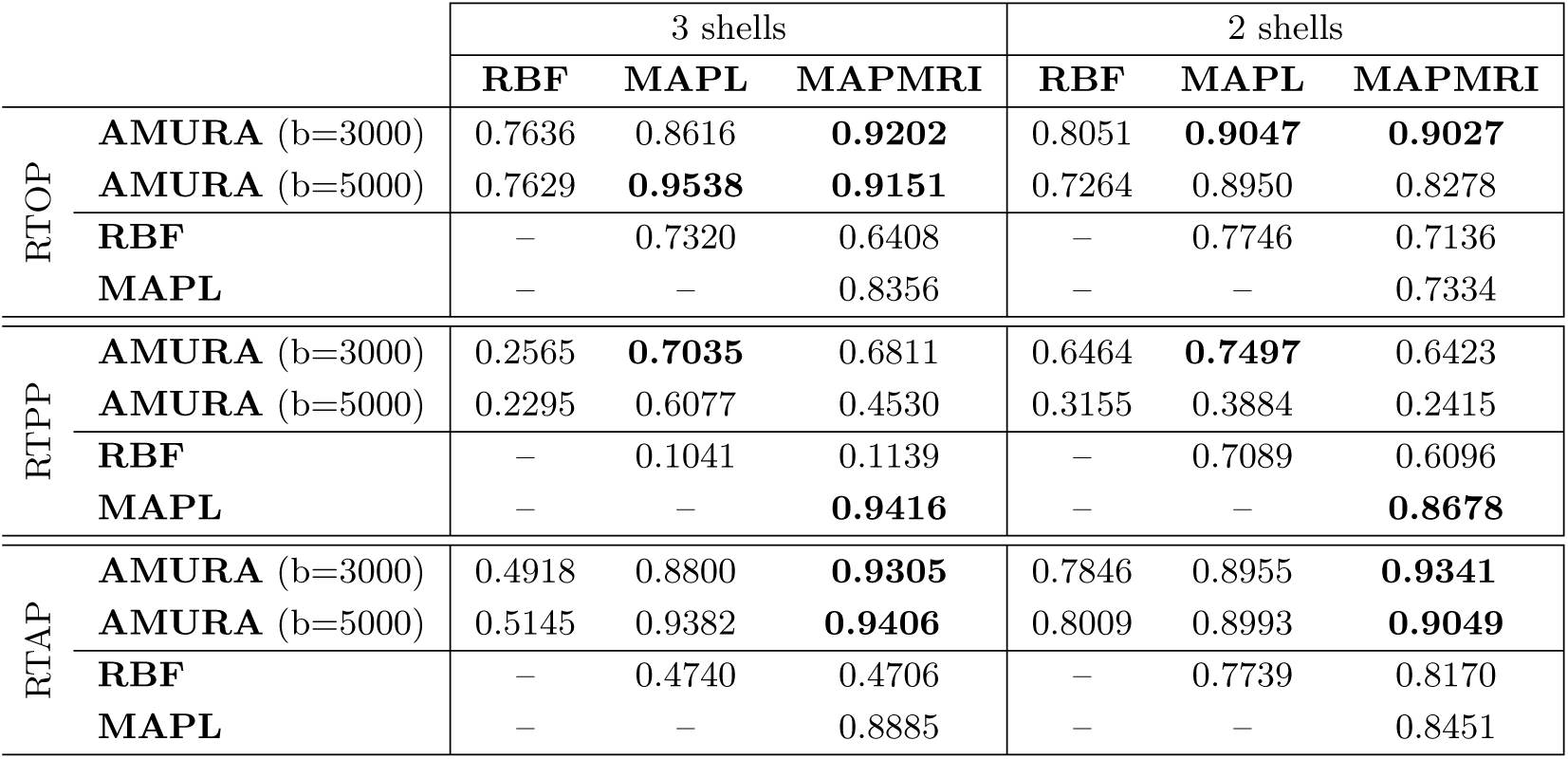
Correlation coefficient between the different methods to estimate RTOP, RTPP and RTAP using 5 different volumes from HCP, the higher the better. The highest values in each case are highlighted in bold face.

**Figure 1:**
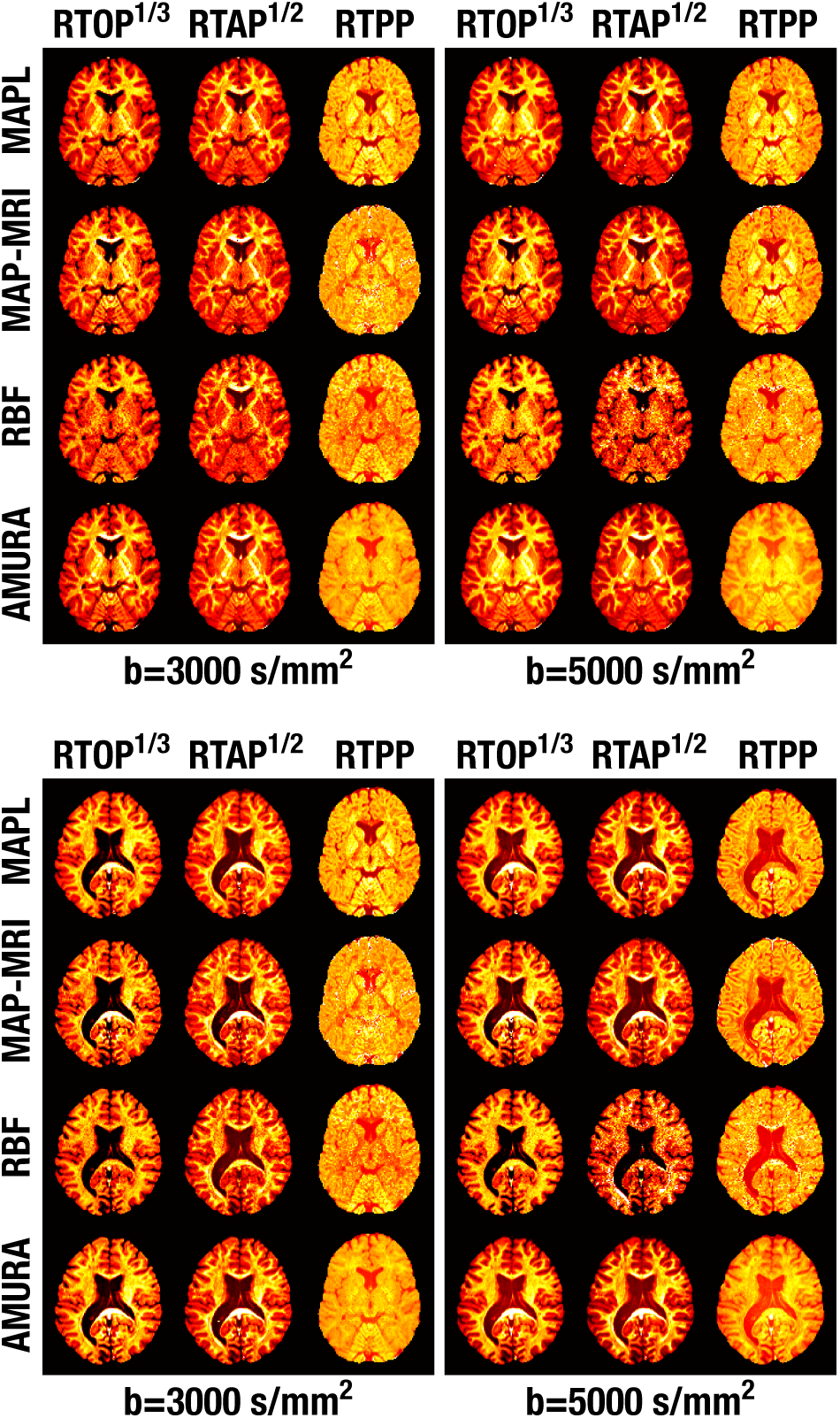
Visual comparison of the q-space measures calculated with different methods. Top: Slice 42 of the MGH1016 volume from HCP; Bottom: Slice 51 of the MGH1018. AMURA is calculated with one single shell. MAPL, MAPMRI and RBF are calculated using 2 and 3 shells (the value of b in the image indicates the highest shell used). For the sake of visual comparison, RTOP and RTAP have been scaled.

Results for RTOP show a strong correlation between the measure estimated with AMURA and the calculation given by the other techniques, particularly those based on MAP. In some cases, the value exceeds the 90% (the correlation with RBF is lower, with its best at 80%). In any case, it is worth noticing that AMURA-RTOP correlates better with MAP-RTOP than RBF-RTOP does, even when RBF is computed from 3 shells (left column) and AMURA is using as few as 64 gradients (b=3000) or 128 gradients (b=5000) in one single shell.

On the contrary, the correlation between AMURA-RTPP and MAP-derived RTPP falls below 75% or 68%, probably meaning more information is lost in this case with the use of one singe shell. Once again, the correlation with RBF-RTPP is weaker (63% at the best). Nonetheless, AMURA still takes over the correlation between MAP-related methods and RBF for RTPP, which can be as low as 10% for the configuration with 3 shells. Interestingly, the computation of RTPP with 2 shells seems more robust (in the sense it exhibits stronger correlations between RBF and MAP) than it is with 3 shells.

The results for RTAP are more coherent: inspecting AMURA-RTAP, the correlations grow up to 90% with MAPL and MAPMRI (but, again, only to 80% with RBF). In the same way as with RTOP and RTPP, the matching between MAP-like and RBF-RTAP is weaker than that between AMURA and MAP-RTAP.

All in all, RTOP and RTAP calculated with AMURA show huge correlations with the measures calculated with state-of-the-art methods, specially with MAPMRI and MAPL, comparable or even above those the other methods yield among them. The situation still holds for RTPP, even when the correlation between different methods is, in absolute terms, significantly weaker for this measure.

Of course, a strong correlation of AMURA with MAPL or MAPMRI does not imply *per se* the same capability of AMURA to discriminate different types of microstructure configurations as MAPL or MAPMRI. However, it is indeed an evidence that the measures calculated with AMURA are measuring some diffusion features closely related to those calculated by MAPL, MAPMRI, and (in a lesser way) RBF. The next experiments, where we evaluate the capability of AMURA to discriminate microstructural features, go in depth into this question.

### 4.3. Discriminant analysis with simulated configurations

The second hypothesis to test is that the proposed measures calculated with AMURA are able to detect microstructural differences in the white matter better than standard diffusion measures like the FA, and with comparable discrimination power to multi-shell, EAP-based methods. The first evidence we provide is based on a set of simulated scenarios based on two compartment models with and without free-water compartment (Panagiotaki et al., 2012). The intra-cellular compartment is modeled by the cylinder and the extra-celular is modeled by zeppelin or ball, see Table. 4. In those cases in which *d* = 3000 *µ*m^2^/s, the isotropic ball corresponds to the free water. Five different voxel configurations are considered, with parameters fine tuned so that they produce the same FA (FA=0.6260) at b=1000 s/mm^2^, see Table 5.

**Table 4:**
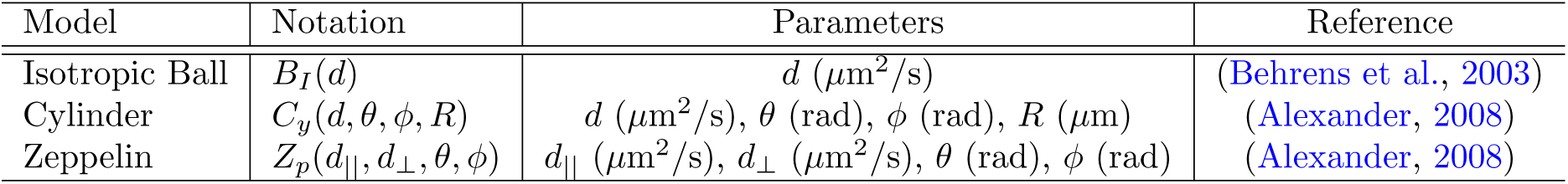
Compartment models used in the simulation.

**Table 5:**
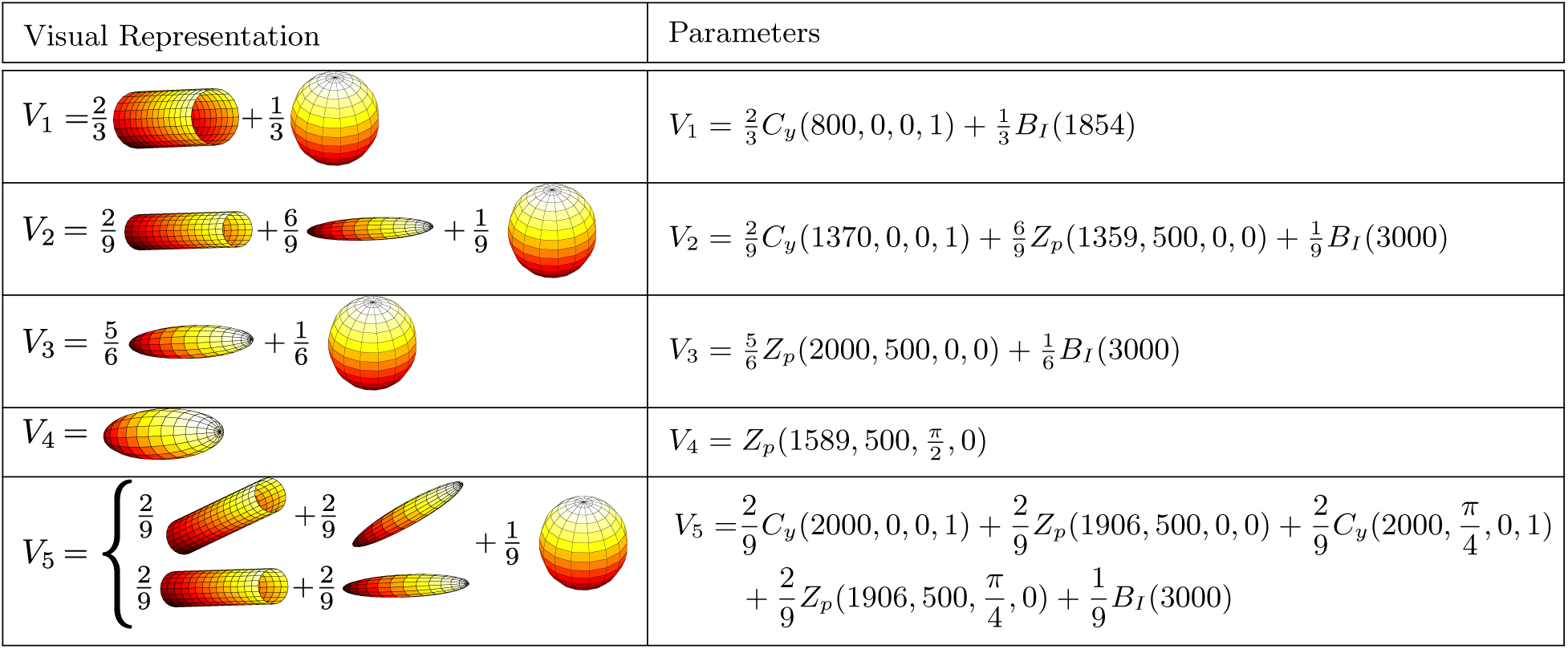
Multicompartment configurations used in the simulation. The distribution and parameters of the models are set so that they have the same FA at *b* = 1000.

In order to simulate the signal, we consider the following sampling scheme: 24 gradient directions uniformly distributed in each shell, 1 baseline, 4 different shells at *b* = [1001, 2019, 3000, 4000] s/mm^2^, Δ = 32.2 ms, *δ* = 27.7 ms, *τ* = Δ *− δ/*3 = 23.0 ms, and *|G|* = [28.3, 40, 48.77, 56.30] mT/m. To statistically analyze the results, we corrupt the data with 30 independent realizations of Rician noise with baseline SNR=40.

RTOP, RTPP, and RTAP are calculated using MAPL (considering the 4 available shells), the DT approach (for each separate shell, see Appendix for formulation) and AMURA (for each separated shell). We have selected MAPL for being the method with the higher correlation results in the previous experiments. In addition, FA value at b=1000 s/mm^2^ is also calculated. In order to test the ability to discriminate the different configurations, a multiple comparison test is carried out over each of the measures in Table 6.

**Table 6:**
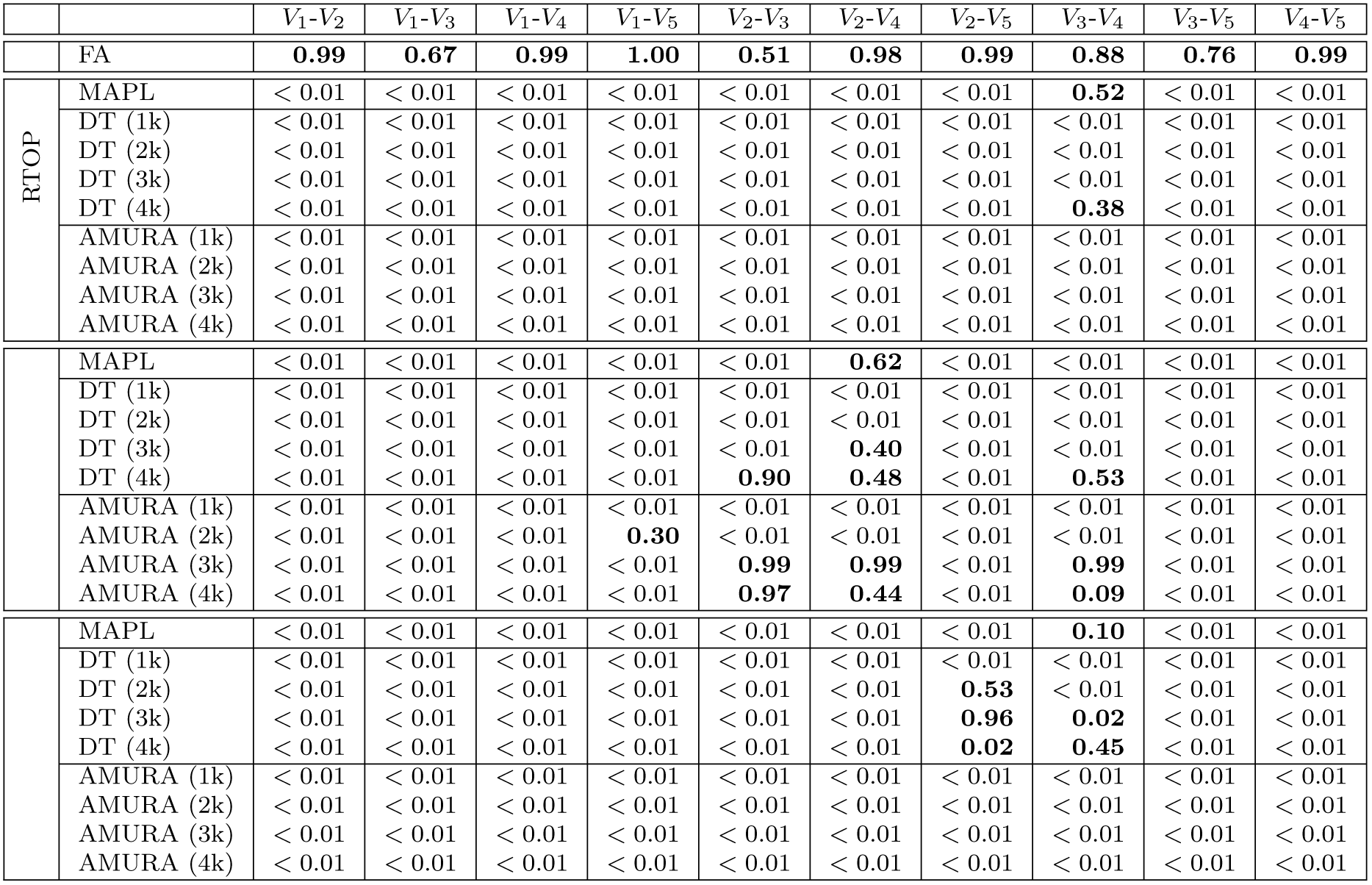
Multiple compare test between 4 voxels with different microstructure. 30 realizations and SNR=40 in the baseline are considered. Differences with statistical significance under 99% are highlighted.

In none of the cases the FA is able to discriminate between voxels, as expected, due to the design of the experiment. RTOP and RTAP calculated with MAPL are both able to tell apart most of the configurations, except Voxels 3 and 4, probably due to some smoothing effect in the estimation of the EAP. DT shows a great discrimination for RTOP, even over MAPL, and slightly worse for RTAP. Some voxels are confused for high b-values, due to the effect of noise over the estimation. Surprisingly, RTOP and RTAP with AMURA are able to discriminate between the different configurations in all of the cases. Note that, even at b=1000, RTOP and RTAP with AMURA are able to discriminate the different values.

On the other hand, RTPP is the index showing the poorer results, for DT and for AMURA. MAPL is only unable to detect 1 configuration, while DT and AMURA work slightly better for low to medium b-values. However, as the b-value grows, their ability to discriminate between voxels highly decreases. This way, for b=4000, both methods are not able to discriminate among 3 different configurations pairs.

### 4.4. Validation with clinical data

The previous simulation is now complemented with the inspection of clinical data, for which we use the PPD database. Though PD is known to affect the substantia nigra or the gray matter more than the *standard* white matter tracts commonly studied in group-wise analyses based on DMRI, significant differences have also been reported in several white matter regions such as the corpus callosum, the corticospinal tract, or the fornix (Atkinson-Clement et al., 2017). Accordingly, we have focused on commonly-studied white matter tracts. To that end, each volume in the data set has been segmented based on the ENIGMA-DTI template^4^ (Jahanshad et al., 2013) and the JHU WM atlas (Mori et al., 2005) as follows:

1. The FA is calculated as a reference value using MRTRIX^5^ with the data at *b*=1000 s/mm^2^.
2. Such map is registered against the ENIGMA-DTI FA template using deformable image registration based on the local cross-correlation between the images (Tristan-Vega et al., 2008). This way we can cope with changes in contrast and illumination produced by differences in the acquisition protocols.
3. Using the JHU WM atlas, 48 different white matter regions of interest are identified and segmented.
4. Given the deformation field provided by the registration algorithm, the segmentations of the ENIGMA-DTI template are back-projected onto the image space of each subject in the PDD. Working on the original image space avoids interpolation artifacts as well as side effects induced by the higher resolution of the ENIGMA-DTI template as compared to the PDD.
5. The ENIGMA-DTI template comprises segmentations of both the whole white matter tracts and their FA skeletons. The back-projection procedure is repeated for both segmentations, so that both a full segmentation of each tract and its pseudo-skeleton (central core) is available in the original image space as shown in Fig 2.
6. Outliers are removed from the segmentations by eliminating those voxels with abnormal values (i.e. values outside the range [0, 1]) of the FA and “Westin’s scalars”, *C*_*p*_, *C*_*l*_, *C*_*s*_ (Westin et al., 2002).

**Figure 2:**
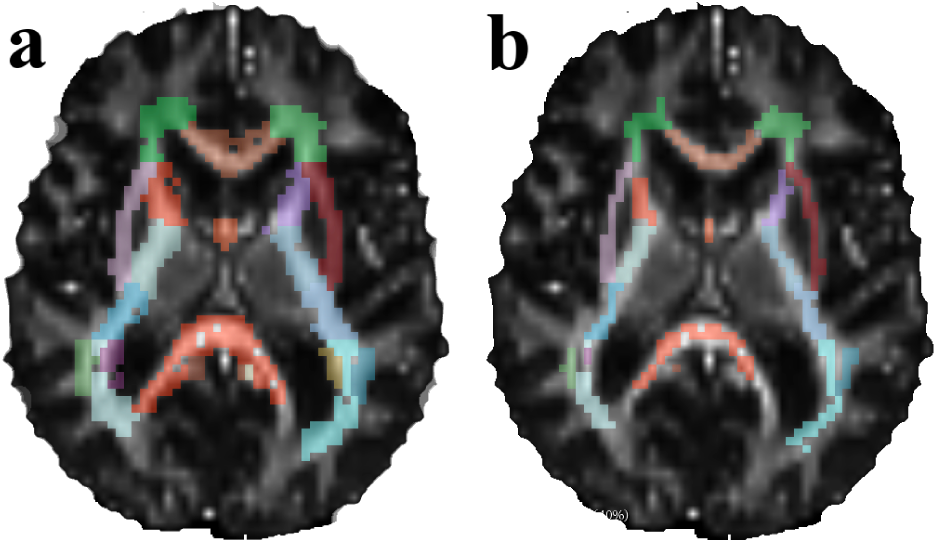
Registration-based segmentation of mainstream white-matter tracts of a given control of the Parkinson data set in its own image space (axial slice), both the whole tracts (a) and their pseudo-skeletons (b).

All the related measures (RTAP, RTOP, and RTPP), computed with either MAPL (the two shells are used), AMURA (at b=2,500 s/mm^2^), or the tensor model (at b=2,500 s/mm^2^), are calculated for all subjects and averaged over the pseudo-skeleton segmentations to obtain one single representative per white matter tract (using the whole tract segmentations produces similar outcomes). The FA is also included in the analysis for comparison purposes. Among the 48 tracts segmented in the JHU WM, we have found statistically relevant differences mainly at the corpus callosum, which is in agreement with the related literature (Atkinson-Clement et al., 2017). Table 7 shows the results for two-sample, pooled variance *t*-tests over Gaussian-corrected data between controls and patients for each of the measures considered and at each of the three sections of the corpus callosum segmented in the JHU WM (genu –GCC–, body –BCC–, and splenium –SCC–). As it can be seen, RTPP-related measures result in discriminant markers for this particular problem at the genu and the splenium of the corpus callosum. Remarkably, the raw FA is only able to find differences at the splenium, meanwhile RTAP and RTOP are unable to plot significant differences in a consistent way.

**Table 7:**
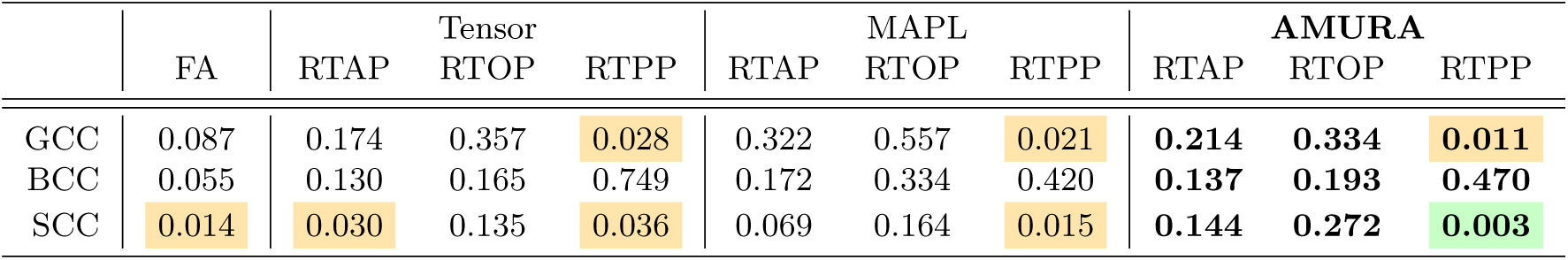
Two-sample, pooled variance, Gaussian corrected *t*-tests for each measure and at each section of the corpus callosum: GCC (genu), BCC (body), and SCC (splenium). The *p*-values represent the probability that the averaged values of the corresponding measure computed over the pseudo-skeleton of the corresponding tract have identical means for both controls and patients. Differences with statistical significance above 99% are highlighted in green, and those with significance over 95% are highlighted in amber.

To further understand why RTPP consistently finds significant differences, and how this is related to the information it measures, its actual distribution (PDF) inside each segment (GCC, BCC, SCC) is estimated by using Parzen windowing. Fig. 3 shows that AMURA-RTPP is able to consistently distinguish between two different populations within each region of the corpus callosum. Meanwhile these two groups are also discriminated at the genu by the other approaches, this is not the case at the body and, above all, at the splenium, where even the MAPL-RTPP fails to find the valley between the two populations. Specifically, statistically significant differences between controls and patients appear wherever there is a change in the relative distribution of voxels between the two populations, i.e., at both the genu and the splenium. This provided, and anytime the separation between the two populations can be easily identified at AMURA-RTPP=13.5 mm^*−*1^, the segmentation of the corpus callosum depending on its RTPP is straightforward by thresholding. Such processing has been applied to each subject in the database (both controls and patients), and the resulting segmentations have been projected onto the image space of the ENIGMA template to compute the average segmentation shown in Fig. 4: The two populations identified by AMURA-RTPP correspond to a clean segmentation of the corpus callosum distinguishing between its lowermost (closer to the cerebrospinal fluid) and its uppermost (closer to the cingulum) sections, so that we can reasonably argue that AMURA-RTPP is actually able to discern microstructural properties that remain hidden with DT-related measures (see Fig. 3).

**Figure 3:**
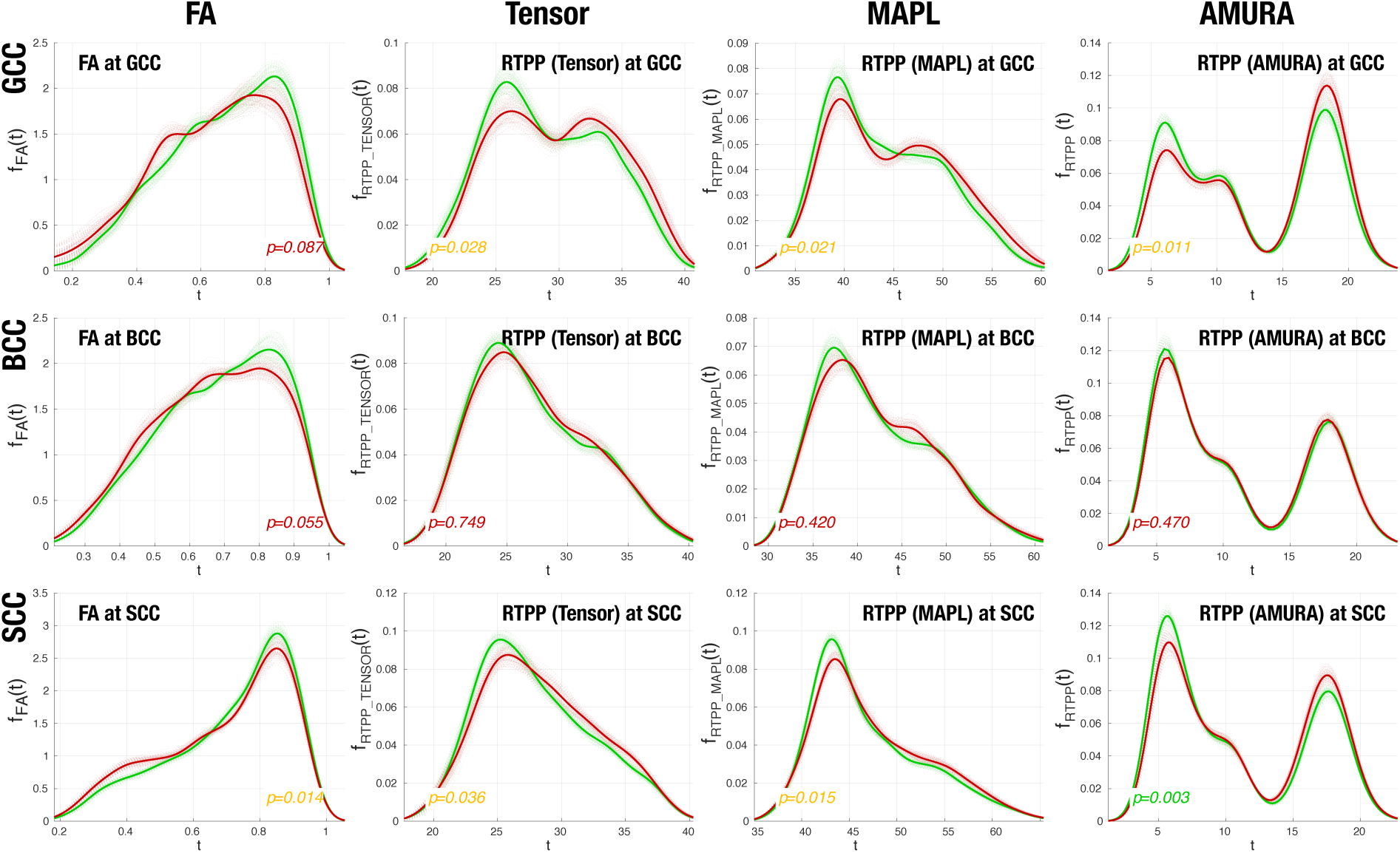
PDFs of each RTPP-like measure (plus the FA) and within each segment of the corpus callosum estimated by Parzen windowing. Only those voxels inside the pseudo-skeleton segmentations are included. Patients (red) and controls (green) are processed independently. Dashed lines correspond to 100 bootstrap iterations accounting for 15 random controls/patients each. Solid lines correspond to the overall, non-bootstrapped PDF in each case. The *p*-values are referred to *t*-tests run over the mean values of each measure over the pseudo-skeleton of each tract, as in Table 7. Top: genu (GCC); middle: body (BCC); bottom: splenium (SCC). From left to right: FA, RTPP computed from the tensor model, RTPP computed from MAPL, and RTPP computed from AMURA.

**Figure 4:**
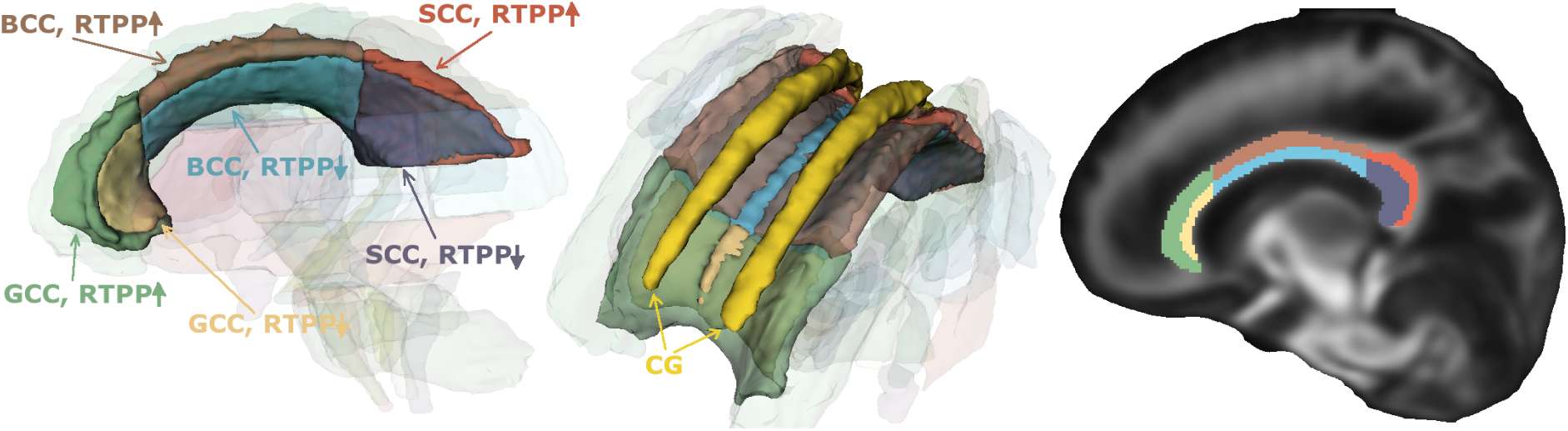
Average segmentation of the corpus callosum in the space of the ENIGMA template by AMURA-RTPP thresholding at 13.5 mm^*−*1^. The cingulum (CG) is also rendered in the 3D view for reference purposes. A sagittal slice of the average FA of the PDD is also shown.

### 4.5. Sensitivity analysis with regard to the acquired shells

The initial assumption for AMURA is that *D*(*θ, ϕ*) does not heavily depend on the b-value. So, next, an experiment aims at checking the sensibility of AMURA to the b-value. One single volume is used for the experiment (MGH 1016), which is further divided in 5 different regions according to their diffusion features. To do that, the FA is first calculated to remove those voxels with FA*<* 0.2. The remaining voxels are clustered in 5 different groups with fuzzy c-means, being the resulting centroids: *C*_*L*_ = {0.24, 0.36, 0.51, 0.66, 0.86}. Each voxel in the white matter is assigned to one cluster using its FA value and the minimum fuzzy distance. AMURA measures are computed for each shell, and the median value inside each of the five clusters is depicted in Fig. 5. RTOP, RTAP and RTPP show a clear dependence with the b-value. This variation, already predicted in section 3.2, is related to the fact that the diffusion at different b-values can be in fact measuring different underlying effects.

**Figure 5:**
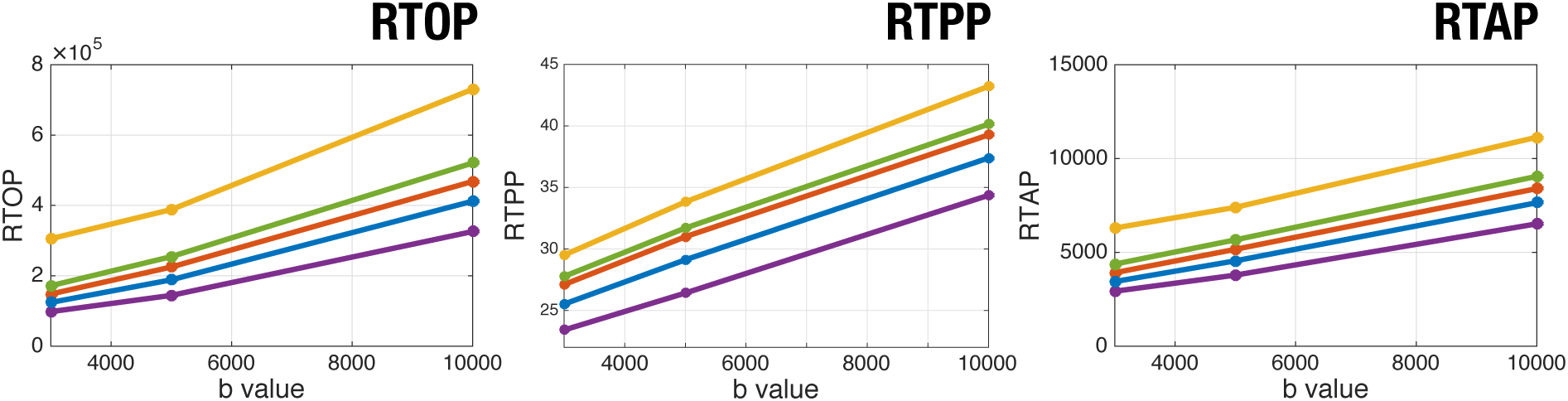
Evolution of AMURA with the b-value, using data from the Human Connectome Project. The volume has been clustered in 5 different sets and the median of each set is shown. Centroids of the data *C*_*L*_ = {0.24, 0.36, 0.51, 0.66, 0.86}.

A similar analysis may be performed over multi-shell state-of-the-art methods: since they depend on the whole set of q-space measures instead of just one shell, they are formerly supposed to be less sensitive to the acquisition protocol. To test this assumption on RBF, MAPL, and MAPMRI, the same five volumes and 3 slices from Table 2 are used. The exact same fuzzy clustering described before is applied now to obtain 5 different classes with centroids *C*_*L*_ = {0.24, 0.33, 0.45, 0.58, 0.76}. Each measure is then computed with either the 2, 3, or 4 innermost shells, and its median depicted in Fig. 6.

**Figure 6:**
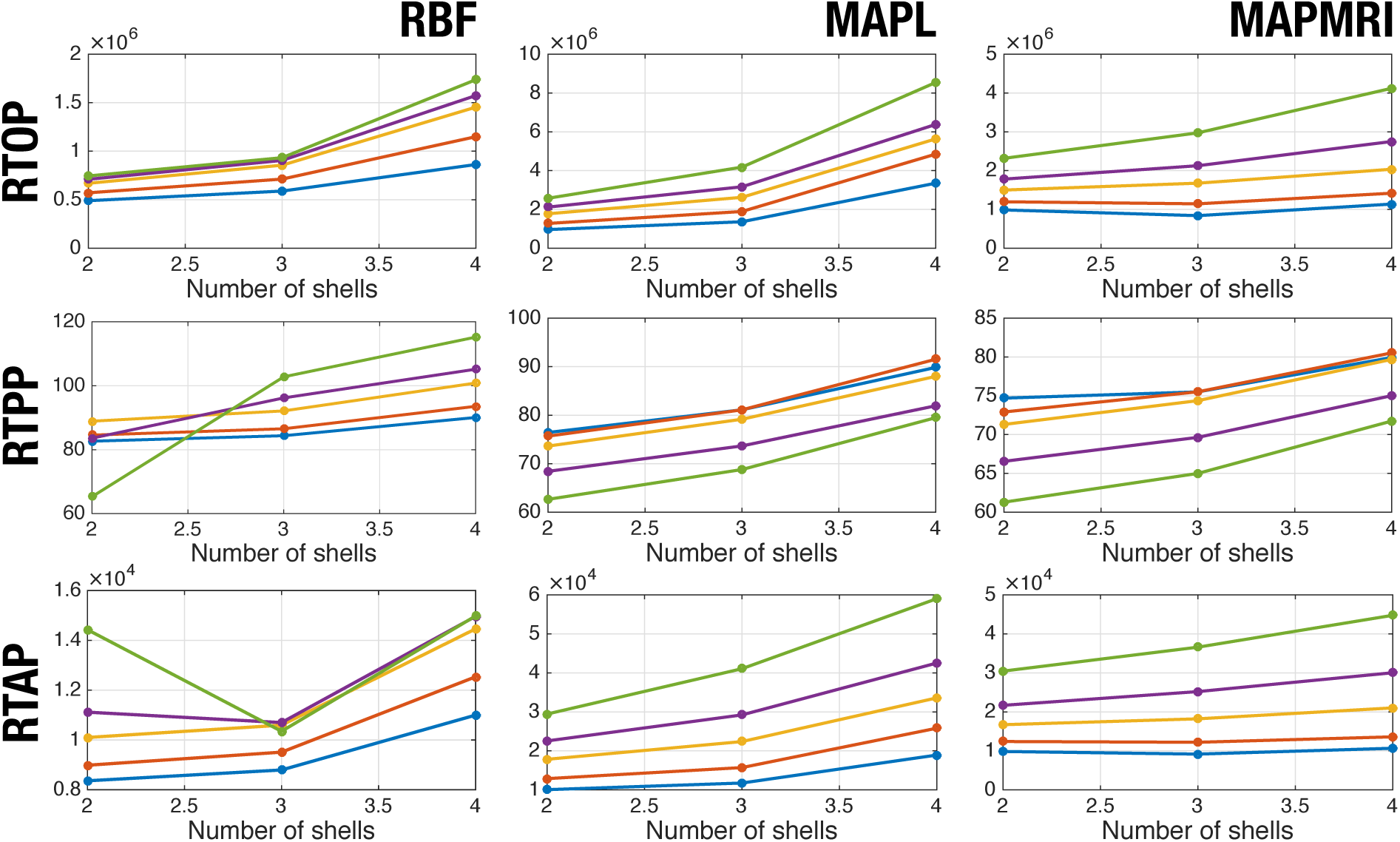
Sensitivity of multi-shell measures to the number of shells used for calculation, using three slices from the 5 volumes in Table 2. The data has been clustered in 5 different sets and the median of each set is shown. Centroids of the data *C*_*L*_ = {0.24, 0.33, 0.45, 0.58, 0.76}.

First thing to notice is that there is an actual dependence of multi-shell measures with the number of shells used for the calculation of the EAP, especially noticeable when the fourth shell is included. For RTOP, even when the output value increases with the number of shells, the relative differences between the 5 segmented regions remain. The same comment holds for RTAP unless the RBF method is considered (which, in the light of this experiment, seems particularly unstable). On the contrary, the evolution of RTPP for each region shows crossings as the number of shells vary, not preserving relative differences.

### 4.6. Execution times

One of the advantages of AMURA is that the SH expansions are derived as linear, regularized LS problems. On the contrary, multi-shell methods avoiding a dense sampling of the q-space often rely on heavily non-linear, sparsity-driven, possibly constrained optimization problems. The linear nature of LS usually yields to well-behaved, stable solutions, meanwhile non-linear optimization usually arises numerical issues.

Besides, the computational load of LS is noticeably more modest (it reduces to invert one single matrix for the whole volume or even the whole cohort), to the point that AMURA can be several orders of magnitude faster than whole EAP-based techniques. To test this extreme, volume MGH 1016 from the HCP is used here to compute diffusion measures on a quad-core Intel(R) Core(TM) i7-6700K 4.00GHz processor under Debian GNU/Linux 8.6 SO. The available code for RBFs^6^ was run under MATLAB 2013b (The MathWorks, Inc., Natick, MA) and the DIPY 0.13.0 library^7^ under Python 3.6.4 (scipy 1.0.0) was used for MAP-MRI and MAPL. AMURA is implemented in MATLAB without multi-threading. The results are reported in Table 8.

**Table 8:** Estimated execution times for the calculation of the measures for the different methods for a single volume (HCP MGH 1016).

| Execution times |  |  |  |
| --- | --- | --- | --- |
| Method | Two shells | Three shells | Four shells |
| RBFs | 118h 10min | 332h 40min | 577h 12min |
| MAP-MRI | 13h 43min | 13h 46min | 16h 20min |
| MAPL | 2h 11min | 2h 14min | 2h 22min |
| AMURA | **6min 41s** | **7min 17s** | **8min 28s** |

Though raw execution times are an ambiguous performance index (they can be dramatically improved, for example, via GPU acceleration), they give a reasonable idea of the complexity of each method. Note the reported times for most of the methods make them unfeasible to be used on practical studies. In the case of RBF, they range from 5 to 24 days per volume, something that goes beyond the capability of clinical groups. Even in the best of the cases, MAPL is 17 times slower than AMURA. In all the cases, most of the time is spent in calculating the EAP. In MAPL, for instance, only 0.6% of the calculation time corresponds to the measures, while the remaining 99.4% is spent in estimating the EAP. In the case of AMURA, 50% of the execution time corresponds to RTAP, since the estimation of the ODF is the most expensive operation, followed by RTPP, which takes 40% of the time. RTOP is the fastest measure, since it takes only 16s, 29s and 54s for the different shells.

## 5. Discussion and conclusions

AMURA-driven measures do not intend to estimate the exact same numeric parameters as EAP-based methods compute. On the contrary, their aim is inferring microstructural information related to, and with comparable discrimination power as, that revealed by MAPMRI, MAPL, or RBF. Fig. 1 and Table 3 evidence the anatomical consistency of AMURA, both visually and numerically. Table 6 and Fig. 3, at least for AMURA-RTPP, confirm its discriminant power over traditional DT markers.

With regard to the first issue, i.e. anatomical consistency, EAP-based measures explicitly account for the radial behavior of the diffusion signal, which is actually sampled. With AMURA, the radial behavior is not sampled but modeled as a mono-exponential decay. The hypothesis leading to the computation of the whole EAP should be, therefore, that the study of the whole EAP provides more specific/sensitive measures, i.e., there is certain anatomical information encoded in the radial behavior of the EAP that would remain hidden with AMURA. However, Table 3 highlights this is not always the case: different EAP-methods bring in less consistent results among them than some of them exhibit with AMURA for analogous measures (RTOP, RTPP, or RTAP). Paradoxically, the similarity between RBF and MAP-like methods even worsens as new shells with higher b-values are introduced.

As a first attempt to explain this behavior, we may recall that the measures computed are merely scalars, i.e., the complex information gathered in the whole 3-D domain of the EAP is somehow collapsed to one single number: the RTOP, for instance, is the value of the EAP at a single point (zero), which corresponds to the integration of the diffusion signal in the whole q-space, in a way that most of the information is lost in the average.

However, the averaging process behind the scalar measures does not explain why the corresponding outputs obtained from the different EAP-based methods do not converge to analogous values, or why the model-constrained AMURA measures seem to mimic MAPL values better than model-free, EAP-based MAPMRI and RBF in Table 3. Moreover, as the number of shells and the number of samples per shell increase, MAPL, MAPMRI, and RBF would be expected to converge to exactly the same values, since all of them estimate the same mathematical entity (the EAP) and all of them use the same mathematical description of the related measures (RTOP, RTPP, and RTAP). The experiments here reported show this is not always the case and, surprisingly, MAPMRI and RBF tend to diverge from MAPL more than AMURA does.

Obviously, the mono-exponential model introduces a non-negligible error in the estimated measures. But the estimation of the EAP is by no means free of certain issues that compromise its accuracy: first, the EAP is usually represented as a superposition of functions selected from a basis or frame where the EAP is assumed to be sparse, which is only a rough approximation; second, the estimation is usually grounded on non-linear, iterative procedures, whose numerical stability is not always guaranteed and whose actual convergence is often conditioned by computational time restrictions; third, EAP estimation requires probing very strong diffusion gradients that drastically worsen the SNR, which may have an uncertain impact depending on the optimization method to be used; fourth, an additional side effect of the use of strong diffusion gradients is that the linear Fourier transform relation between the EAP and the diffusion signal, which is the keystone of all EAP-based methods, may no longer hold with accuracy due to non-linearity, diffraction, and/or non-negligible diffusion during the application of pulsed gradients in a time *δ* (see Fig. 6, where including the fourth shell in the estimation heavily increases all measures for MAPL; this may suggest the Fourier model has been compromised at this point).

The combination of these four factors (and possibly others) may affect each EAP-based method in very different ways, and they could even represent a larger error than that introduced by the mono-exponential model. This could possibly explain the discrepancies between the measures computed with any of the three EAP-based methods, especially the higher deviations of RBF when 3 shells (instead of 2) are used in Table 3. For instance, the very same voxel may show RTOP values as different as 1.2 × 10^5^ with RBF, 1.0 × 10^5^ with MAPMRI, or 1.0 × 10^5^ with MAPL. Of course, AMURA does not get rid of this issue, and the corresponding value may range 2.0 × 10^5^. This scale-shift may be confirmed for RTPP in Fig. 3, too. But, once again, the goal of AMURA is not estimating the exact same values as EAP-based methods: a shifted (level/contrast changed) version of a given measure will have exactly the same discriminant power as its former version, and therefore it will be equally valuable. Going back to Fig. 6, EAP-based measures do not always respect this principle: depending on the number of shells used, certain measures (RTPP and RTAP) are assigned very different, non-consistent relative values among the anatomies compared. This becomes especially annoying with RBF, for which crossings of values for RTPP and RTAP appear as new shells are incorporated. MAPL and MAPMRI prove themselves more stable but, even when the values are not crossing, some of them converge. Finally, RTPP seems more sensitive to this problem: as opposed to RTOP and RTAP, which represent averaged magnitudes over the unit sphere, RTPP relies only on the main diffusion direction, making its computation less robust.

Once the consistency of the measures computed with each method has been thoroughly discussed, the big deal is their power to resolve microstructural features beyond the capabilities of conventional DT-MRI. Table 6 suggests that both RTOP and RTAP have a great potential in this sense, or at least comparable to the potential of MAPL. All the configurations distinguished by MAPL with RTOP and RTAP are satisfactorily resolved by AMURA.

The discussion changes when referring to RTPP: while for low-to-medium b-values AMURA is still able to find most of the differences, its ability to discriminate between voxels dramatically decreases as the b-value increases (and, as a consequence, the SNR worsens). This way, 5 configurations-pairs are lost out of 10 for b=6000 s/mm^2^. Remarkably, MAPL loses 3 out of these 10 pairs even when it is using 6 shells. As stated before, AMURA-RTPP is computed from just one single diffusion direction (instead of being an averaged value), hence it is expected to be more prone to noise and estimation artifacts. This is also in good agreement with the poor correlation results reported in Table 3 for this measure.

If we focus now on the experiment in Section 4.4, the discussion seems to change once again when clinical data are involved: while RTOP and RTAP are unable to find statistically significant differences between controls and patients, RTPP does, see Table 7. It is important to stress here that the aim of the experiment is not demonstrating the clinical usefulness of AMURA in the particular case of PD, but testing its capability to describe microstructural features. In other words, the fact that RTOP and RTAP are not able to find significant differences between controls and PD patients only means that the microstructural properties they describe do not seem to be altered by this particular pathology and/or this particular dataset.

Nonetheless, even when RTPP seemed less useful than RTOP and RTAP in the light of Tables 3 and 6, it provides highly valuable information for the particular case of PD. As opposed to the previously discussed cases, where the voxel-wise RTPP was studied, Table 7 represents statistical tests run on average RTPP values computed over the whole segmentation of the corpus callosum, which should notably improve its robustness to noise and estimation artifacts. On the other hand, Fig. 3 suggests the capability of RTPP to tell controls from patients comes from the change in the relative populations of two well-defined regions that are not always described by the DT-derived measures or even the MAPL-derived. Of course, the PDD is not particularly well suited for MAPL estimation since only 2 shells at relatively low b-values are available, so the latter comparison is not completely fair.

Paying attention to Fig. 4, the two populations distinguished by AMURA-RTPP become evident: in the outermost region, the corpus callosum is interleaved with the cingulum, so that restricted diffusion prevails, the maximum diffusivity decreases and the RTPP increases (lobes at the right of the valley in Fig. 3, rightmost column). In the innermost part, on the contrary, the corpus callosum is closer to the CSF and non-restricted diffusion takes a more relevant role: the maximum diffusivity increases and, as a consequence, the RTPP decreases (lobes at the left of the valley in Fig. 3). Comparing this reasoning with the experiment in Section 4.3, RTPP seems to fail in the description of subtle microstructural changes, but it could have higher sensibility/specificity to discriminate scenarios with different amounts of free water.

All in all, we can conclude that AMURA shows a better ability to discriminate among different microstructural configurations for very different scenarios than conventional DT techniques. At the same time, such ability compares to that of sophisticated techniques based on multi-shell acquisitions and large data sets. Though AMURA would obviously benefit from a larger number of samples, its main advantage is that it remains compatible with nowadays standard acquisition protocols (with as few as 64 gradient directions): it is a common practice acquiring two shells (b=[1000, 3000] s/mm^2^, for instance) to estimate classical diffusion parameters, like the FA and MD, but also advanced models (DKI, HARDI, CHARMED, etcetera). AMURA allows to add RTOP, RTPP, and RTAP to the pool with no additional effort and without changing the acquisition protocol, in the same way it was done in Section 4.4 for the already existing PDD.

Moreover, since AMURA avoids the estimation of the actual EAP, the computation of its related measures may be done in a fast and robust way, i.e., without imposing a computational burden to the standard protocols: some of the experiments in the present paper report an acceleration about three orders of magnitude (10^3^) compared to EAP-based measures, see Table 8. A whole volume can be processed in 6 to 8 minutes, so that a clinical study with 200 different subjects could be finished in 26 hours. The same cohort would take 4808 days (RBF), 135 days (MAPMRI), or 19 days (MAPL), which obviously limits the applicability of these methods. The computational simplicity of AMURA, however, does not only imply faster execution times, but also more robust estimations due to its closed-form, mainly linear nature. As opposed, EAP-based techniques usually estimate the whole EAP from multi-shell samplings based on non-linear methods, which, as discussed above, lead to high discrepancies in the output measures.

On the other side of the coin, the major drawback behind AMURA is the explicit assumption of a specific radial behavior for the diffusion, which cannot fit the whole q-space. As a consequence, the selection of the b-value may change the values of the measures. However, as we have shown, this variability with the b-value can also be found in other state-of-the-art methods (see Fig 6), whose results depend on the shells used for the estimation of the EAP. Based on this result, we have to consider our measures as *apparent values* at a specific b-value. This implies the results of clinical trials could be compared against each other only if the same b-value is preserved across the studies. This is by no means something new to diffusion imaging: it is well-known that a change in the acquisition parameters (number of gradients, b-value, resolution, scanner vendor, etcetera) seriously affects scalar measures like the FA or the MD (Aja-Fernández et al., 2018; Barrio-Arranz et al., 2015).

## Acknowledgments

This work was supported by Ministerio de Ciencia e Innovación of Spain with research grants RTI2018-094569-B-I00 and PRX18/00253 (Estancias de profesores e investigadores senior en centros extranjeros). Maryam Afzali is supported by a Wellcome Trust grant (096646/Z/11/Z). Tomasz Pieciak acknowledges National Science Centre (Poland) for funding resource (2015/19/N/ST7/01204)

The authors acknowledge Lipeng Ning, Carl-Fredrik Westin and Yogesh Rathi from Brigham and Women’s Hospital, Harvard Medical School for sharing the source code of directional RBFs and their outstanding assistance. The authors thank the contributors of DIPY project (http://nipy.org/dipy/) for providing the MAP-MRI basis implementation.

Data collection and sharing for this project was provided by (1) the *Human Connectome Project* (HCP; Principal Investigators: Bruce Rosen, M.D., Ph.D., Arthur W. Toga, Ph.D., Van J. Weeden, MD). HCP funding was provided by the National Institute of Dental and Craniofacial Research (NIDCR), the National Institute of Mental Health (NIMH), and the National Institute of Neurological Disorders and Stroke (NINDS). HCP data are disseminated by the Laboratory of Neuro Imaging at the University of Southern California; (2) the *High-quality diffusion-weighted imaging of Parkinson’s disease* data base, Cyclotron Research Centre, University of Liége.

## A. Calculation of the structural measures using the diffusion tensor

If a Gaussian diffusion propagator is assumed, *P*(**R**) is a mixture of independent and (nearly) identically distributed bounded cylinder statistics and, by virtue of the central limit theorem, their superposition is Gaussian distributed. The measured signal in the q–space is the (inverse) Fourier transform of the PDF and it can be expressed as:

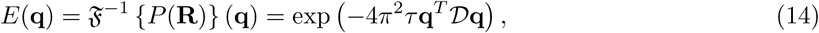

which represents the well-known Stejskal–Tanner equation (Stejskal and Tanner, 1965). The diffusion tensor *𝒟* is the anisotropic covariance matrix of the Gaussian PDF *P* (**R**), and therefore is a symmetric, positive–definite matrix, with positive eigenvalues and orthonormal eigenvectors. If we use this model to estimate the measures, we obtain:

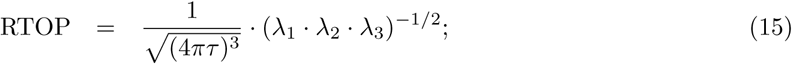

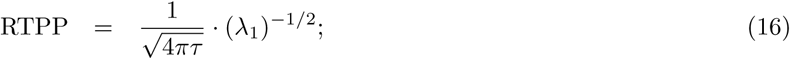

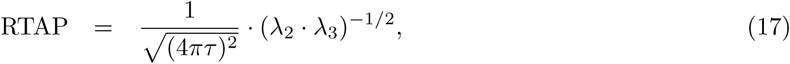

where *λ*_1_ ≥ *λ*_2_ ≥ *λ*_3_ are the three real, non-negative eigenvalues of *𝒟*.

## Footnotes

1 Data obtained from the Human Connectome Project (HCP) database (https://ida.loni.usc.edu/login.jsp). The HCP project (Principal Investigators: Bruce Rosen, M.D., Ph.D., Martinos Center at Massachusetts Gen eral Hospital; Arthur W. Toga, Ph.D., University of Southern California, Van J. Weeden, MD, Martinos Center at Massachusetts General Hospital) is supported by the National Institute of Dental and Craniofacial Research (NIDCR), the National Institute of Mental Health (NIMH) and the National Institute of Neurological Disorders and Stroke (NINDS). HCP is the result of efforts of co-investigators from the University of Southern California, Martinos Center for Biomedical Imaging at Massachusetts General Hospital (MGH), Washington University, and the University of Minnesota.

2 The SNR of each of the individual baselines is high enough to make a Gaussian approximation feasible with a small error. Under this approximation we can assure that the average operator provides an unbiased output image (Aja-Fernández and Vegas-Sánchez-Ferrero, 2016).

3 Acquired at the Cyclotron Research Centre, University of Liége. Available: https://www.nitrc.org/frs/?group_id=835.

4 ENIGMA project web page: http://enigma.ini.usc.edu/. Template data and processing protocols for DTI: https://www.nitrc.org/projects/enigma_dti.

5 Available at: http://www.mrtrix.org.

6 https://github.com/LipengNing/RBF-Propagator.

7 http://nipy.org/dipy.

